# Intra-genomic rDNA gene variability of Nassellaria and Spumellaria (Rhizaria, Radiolaria) assessed by Sanger, MinION and Illumina sequencing

**DOI:** 10.1101/2021.10.05.463214

**Authors:** Miguel M. Sandin, Sarah Romac, Fabrice Not

## Abstract

Ribosomal DNA (rDNA) genes are known to be valuable markers for the barcoding of eukaryotic life and its phylogenetic classification at various taxonomic levels. The large scale exploration of environmental microbial diversity through metabarcoding approaches have been focused mainly on the V4 and V9 regions of the 18S rDNA gene. The accurate interpretation of such environmental surveys is hampered by technical (e.g., PCR and sequencing errors) and biological biases (e.g., intra-genomic variability). Here we explored the intra-genomic diversity of Nassellaria and Spumellaria specimens (Radiolaria) by comparing Sanger sequencing with Illumina and Oxford Nanopore Technologies (MinION). Our analysis determined that intra-genomic variability of Nassellaria and Spumellaria is generally low, yet some Spumellaria specimens showed two different copies of the V4 with <97% similarity. From the different sequencing methods, Illumina showed the highest number of contaminations (i.e., environmental DNA, cross-contamination, tag-jumping), revealed by its high sequencing depth; and MinION showed the highest sequencing rate error (~14%). Yet the long reads produced by MinION (~2900 bp) allowed accurate phylogenetic reconstruction studies. These results highlight the requirement for a careful interpretation of Illumina based metabarcoding studies, in particular regarding low abundant amplicons, and open future perspectives towards full environmental rDNA metabarcoding surveys.

## Introduction

Ribosomal DNA (rDNA) genes are known to be valuable taxonomic markers for the barcoding of eukaryotic life and its phylogenetic classification at different levels; mainly due to its intra-genomic tandem repeated structure, the presence of conserved and variable regions and its appearance in all eukaryotes (Pawlowski et al., 2012; del Campo et al., 2018). The 18S rDNA gene, coding for the small sub-unit of the ribosome, has been widely used in molecular environmental surveys, especially the short hyper-variable regions V4 and V9, thanks to the extensive occurrence in public databases and the availability of generalist primers flanking their sides (Amaral-zettler et al., 2009; Stoeck et al., 2010). The advent of High-Throughput Sequencing (HTS) techniques has allowed the massive sequencing of molecular environmental diversity supporting its exploration through a metabarcoding approach (de Vargas et al., 2015; Massana et al., 2015; Forster et al., 2016; Pernice et al., 2016). The large amount of reads (or amplicons) generated by HTS is normally classified in Operational Taxonomic Units (OTUs) based on arbitrary similarity thresholds. OTUs are not only used to identify taxonomic entities, but also to describe community structure (Blaxter et al., 2005). The increasing use of the HTS, have led to the development of different clustering methods resulting in finer-scale OTUs that focus in single nucleotide differences (Mahé et al., 2015) or in the correction of sequencing errors based in the error rate entropy (so-called Amplicon Sequence Variants or ASVs; Callahan et al., 2016).

HTS produce a vast amount of reads carrying errors that are difficult to distinguish from real biological variation, that is considered as a main factor inflating diversity (Kunin et al., 2010). Intra-genomic rDNA Polymorphism and its different copy numbers among taxa can also affect diversity assessments (Gong et al., 2013; Gong and Marchetti, 2019). Other less common causes, yet important, have also been reported as factors inflating diversity estimates, such as lateral gene transfer (Yabuki et al., 2014) or presence of pseudogenes (Thornhill et al., 2007). Considering the very high sequencing depth allowed by the current HTS technologies it is likely that most of this intra-genomic diversity could be sequenced, potentially leading to a large overestimation of the environmental diversity. Several studies have argued that, quantitatively at the community level, the amount of molecular clusters largely exceeds that of morphological counts (Medinger et al., 2010; Bachy et al., 2013; Santoferrara et al., 2016), leading to hypothesis that using HTS approaches scientists are actually measuring species intra-genomic variability (Caron and Hu, 2018). Current HTS technologies used for environmental surveys can sequence short fragments of DNA of about 400 base pairs (bp) only, such as the hyper-variable regions (V4 and V9 most commonly used in protist) of the 18S rDNA gene. Comparing such short hyper-variable regions to, far from exhaustive reference sequences databases (Pitsch et al., 2019) may also contribute to inflating environmental diversity by misidentification of environmental clusters or lack of intra-genomic rDNA variability representation, among other causes.

New sequencing technologies have been developed with the ability of high throughput sequencing longer nucleotide fragments in real time, such as Oxford Nanopore Technologies (ONT) or Pacific Bioscience (PacBio). These sequencing methods have already showed their useful capabilities, for example PacBio have developed a circular consensus sequencing resulting in near-zero error long reads, improving aspects such as genome assembly (Wenger et al., 2019) or even phylogenetic analysis of environmental diversity (Jamy et al., 2019). However, its limited high throughput sequencing ability and its relatively high cost (Goodwin et al., 2016), may affect the sequencing depth at a metabarcoding community level. On the other side, ONT provides a large amount of reads, inexpensively and highly portable with the MinION device (Levy and Myers, 2016). Despite that the error rate of the ONT reads are improving since it was firstly released (for MinION, from ~60% in 2014 to <15% error rate; Rang et al., 2018) it is still a major concern, reaching up to 3-6% of errors in the best case scenarios (Tyler et al., 2018).

In this study, we assess the intra-genomic variability within protists by comparing three different sequencing methods, Sanger, ONT (MinION) and Illumina. We focused our efforts in two groups of Radiolaria, the Nassellaria and Spumellaria (Polycystines). The Radiolaria is an important group of protists in the eukaryotic plankton community contributing for a major fraction of the total reads in environmental molecular surveys (de Vargas et al., 2015; Pernice et al., 2016). However, the number of morpho-species described among Radiolaria taxa (Suzuki and Not, 2015) does not match with the molecular barcodes found in environmental surveys. Nassellaria and Spumellaria environmental diversity lags far behind other radiolarian, despite of possessing the largest morphological diversity described. Recent studies have dwelled on exploring their extant morpho-molecular diversity (Sandin et al., 2019; Sandin et al., 2021) showing their uncharted diversity among Radiolaria. The ecological importance of these groups and their observed low molecular diversity in environmental surveys stresses the need for understanding such differences among molecular markers and morphological diversity.

## Material and Methods

### Single-cell Sampling, isolation and DNA extraction

Plankton samples were collected in the Bay of Villefranche-sur-Mer (France) and in the West Mediterranean Sea (MOOSE-GE 2017 expedition) by plankton nets tows (from 20 to 64 μm mesh size). More information on sampling methodology for specific samples can be found in the RENKAN database (http://abims.sb-roscoff.fr/renkan/). Specimens were individually handpicked with Pasteur pipettes from the plankton community and maintained in 0.2 μm filtered seawater for several hours. They were transferred 3-4 times into new 0.2 μm filtered seawater to allow self-cleaning from debris, particles attached to the cell or prey(s) digestion. By doing so, it is expected to keep only essential entities from the holobiont (the radiolarian host + associated symbionts and bacteria). Live specimens were imaged using an inverted microscope and thereafter transfer into 1.5 ml Eppendorf tubes containing 50 μl of molecular grade absolute ethanol and finally stored at −20 °C until DNA extraction. DNA was extracted independently for each specimen using the MasterPure Complete DNA and RNA purification Kit (Epicentre) following manufacturer’s instructions.

### Amplification and sequencing

A schematic representation of the study design and the different amplification and sequencing steps can be found in Supplementary material Fig S1.

Four holobionts were selected to amplify the full length of the rDNA gene (Supplementary material Fig S2), 2 of them belonging to Spumellaria: Mge17-81 (*Rhizosphaera trigonacantha*, Rhizosphaeroidea) and Mge17-82 (*Spongosphaera streptacantha*, Spongosphaeroidea); and 2 to Nassellaria: Vil325 (*Eucyrtidium cienkowskii*, Eucyrtidioidea) and Vil496 (*Eucyrtidium acuminatum*, Eucyrtidioidea). Each holobiont was PCR-amplified in three different technical replicates with the eukaryotic primers SA (AACCTGGTTGATCCTGCCAGT) / D2C-R (CCTTGGTCCGTGTTTCAAGA) (Medlin et al., 1988; Scholin et al., 1994). Amplifications were performed with the enzyme Phusion® High-Fidelity DNA Polymerase (Finnzymes). The PCR mixture contained 1μL (~0.5ng) of DNA template with 0.35μM of each primer (final concentration), 3% of DMSO and 2X of GC buffer Phusion MasterMix (Finnzymes), in a final volume of 25μL. Amplifications were done in a SIMPLIAMP (Applied Biosystems™) with following the PCR program: initial denaturation step at 98°C for 30 s, 37 amplification cycles of 10 s at 98°C, 30 s at 55°C, 30 s at 72°C and final elongation step at 72°C for 10 minutes. For each of the 12 amplified samples, a 23-base pairs (bp) multiplex identifier or BC sequence was designed and included in one of the primers in order to identify the origin of every single read from the pooled population generated on a single run. Structures of the “BC- primers” were as follows: Primer 1: (5′- Tag + Forward primer -3′); Primer 2: (5′- Reverse primer -3′). To avoid single cell DNA sample contamination, PCR mixtures were carried out under a DNA–free vertical laminar flow hood. Negative and positive controls were added to check if there were no sample contaminations (DNA from operator, reagents or material contamination, etc.). PCR products were then purified using NucleoSpin®Gel and PCR Clean up kit (Macherey-Nagel™). Final elution product was divided in two for sequencing by Sanger (after cloning) and Oxford Nanopore technologies using the MinION device (Laver et al. 2015, Jain et al. 2015).

#### · Sanger sequencing

After purification, PCR amplicons were then cloned using TOPO TA Cloning for sequencing kit (Invitrogen) following manufacturer’s recommendations. After a first step of 6 min ligation into pCR™4-TOPO® vector, ligated PCR products were then transformed into TOP10 Chemically Competent *Escherichia coli* (ampicillin resistant) by heat-shock treatment at 42°C for 30 s. Cells were then incubated 2h at 37°C with rotation tray at 200 rpm with an enrichment medium. Afterwards, 80 μL of each transformant was spread on selective solid LB-agar medium (Sigma) with 50 μg/mL of ampicillin and incubated overnight at 37°C. On the following day, presence of the insert was verified in obtained colonies by PCR using the primers M13 forward and M13 reverse complementary to cloning site flanking sequences. A total of 24 clones (4 holobionts x 3 replicates x 24 clones = three 96-well plates) from each amplicon were amplified directly with GoTaq polymerase (Promega, Lyon France) in a 25 μL reaction volume using the following PCR parameters: 10 min at 95 °C, 35 amplification cycles of 30 s of denaturation at 95 °C, 30s annealing at 56°C and 1 min extension at 72 °C, with a final elongation 10 min at 72 °C. PCR products were then purified using ExoStar (Illustra™ Exostar™ 1_Step, GE Healthcare Bio-Sciences Corp.). Final PCR products were sent to Macrogen Europe for sequencing using the primers SA (AACCTGGTTGATCCTGCCAGT; Medlin et al., 1988), S69f (AAHCTYAAAGGAAHTGACGG), D1Rf (ACCCGCTGAATTTAAGCATA; Scholin et al., 1994).

#### · MinION library preparation and sequencing

Each PCR product was pooled together in equimolar conditions to get 200 ng of final amplicon concentration in 45μL. Quality, size, and concentration of PCR products were assessed on a Bioanalyzer 2100 (Agilent) using a DNA1000 Labchip and on a Qbit fluorometer using a Qbit HS DNA quantification kit (Invitrogen). For this study, we optimized a protocol using Oxford Nanopore Technology (ONT) 1D^2 sequencing chemistry combined with Flowcell R 9.5.

Library preparation: DNA products (45 μL amplicon) were treated by an end-repair/dA tailing using NEBNext® Ultra™ II End Repair/dA-Tailing Module (New England Biolabs) and NEB Blunt/TA Ligase Master Mix (New England Biolabs), and D2 Adapter. Samples were washed using AMPure beads (Agencourt) and 70% EtOH and re-suspended in 45 μL water. A first step of adapter ligation was conducted with 25 μL NEB Blunt/TA Ligase Master Mix (New England Biolabs), and 2.5μL 1D^2^ Adapter, and then carried out with 5μL BAM and 50μL of Blunt/TA ligase. A second washing step was done using AMPure beads (Agencourt) and 70% EtOH and re-suspended in 15 μL water of final elution.

Sequencing: Pre sequence-library was prepared following ONT protocols by mixing 12 μL amplicon, 2.5 μL water, 25.5 μL LBB and 35 μL RBF. Flow cell R 9.5 was inserted in the MinION frame and priming port was loaded with 800 μL of priming mix (576 μL RBF and 624 μL water) with SpotON cover closed. We additionally loaded 200 μL of priming mix into the flowcell via the priming port, avoiding the introduction of air bubbles. 75μL of sample library was then added to SpotOn port via dropwise fashion. Finally, we covered SpotOn and priming ports, close the MinION lid and open the MinKNOW GUI software to proceed with sequencing using a half of a total flow cell R 9.5. It is important to note that the flowcell was used during its last 28h (20h → 48h), after a prior independent experiment (data being analyzed).

#### · Illumina sequencing

In total 8 holobionts were selected to amplify the V4 region (~380pb) of the 18S rDNA gene (Supplementary material Fig S2), 4 belonging to Spumellaria: Mge17-81 (common to previous section), Mge17-82 (common to previous section), Vil480 (*Tetrapyle octacantha*, Pylonioidea), Vil497 (*Arachnospongus varians*, Liosphaeroidea); and 4 to Nassellaria: Mge17-9 (*Extotoxon undulatum*, Artostrobioidea), Mge17-124 (*Carpocanium obliqua*, Carpocaniidae), Vil490 (*Pterocorys* cf. *zanclea*, Pterocorythoidea), Vil496 (common to previous section). Specimen Vil325 (from previous section) had no DNA left for further experiments and therefore was not possible to include in this section. Each holobiont was PCR-amplified with the eukaryotic primers TAR-EukF1 (5’-CCAGCASCYGCGGTAATTCC-3’) / TAR-EukR3 (5’-ACTTTCGTTCTTGATYRA-3’) (Stoeck et al. 2010). The forward primer had a barcode adapted for Illumina sequencing and each holobiont was amplified with 3 different barcode-adapted-primers to get 3 technical sequencing replicates. PCR reactions contained 1x MasterMix GC Phusion High-Fidelity DNA Polymerase (Finnzymes), 0.35μM of each primer, 3% dimethylsulphoxide and 1μL of DNA in a final volume of 25μl. The PCR program had an initial denaturation step at 98°C during 30 s, followed by 15 cycles of 10 s at 98°C, 30 s at 53°C and 30 s at 72°C, then 22 similar cycles but with 48°C annealing temperature, and a final elongation step at 72°C for 10 min. Vil325 gave no positive reactions and therefore was no possible to be included in this step. Polymerase chain reaction, in triplicates, were purified with NucleoSpin Gel and PCR Clean-Up kit (Macherey-Nagel), and quantified with the Quant-It PicoGreen double stranded DNA Assay kit (Invitrogen). About 500 ng of pooled amplicons were sent to Fasteris (https://www.fasteris.com, Switzerland) for Illumina sequencing on a Miseq nano V2 2×250.

Sequencing results from this study along with associated metadata for its analysis and native formats have been deposited in figshare with the following DOI: 10.6084/m9.figshare.16922764.

All scripts used in this study along with the tools for the replication of the analysis are available in github (github.com/MiguelMSandin/IntraGenomic-variability).

### Quality checking and similarity of sequences/amplicons

In order to compare the output of the three different sequencing results, similar and broadly known methods were used to quickly check the quality of the sequences/amplicons obtained. Firstly, raw sequences/amplicons were measured in base pairs (bp) length. Thereafter, sequences/amplicons were compared against reference sequences by local alignment (BLAST; using NCBI as reference sequences) for Sanger and MinION sequencing results and global alignment (vsearch; Rognes et al. 2016; using PR2 v4.14.0 database as reference sequence, Guillou et al. 2013) for Illumina results. This difference in the alignment algorithm used is due to the big differences in the sequence length and the method mostly used for each of the sequencing results. Finally, in order to obtain sequences per holobiont, Sanger reads belonging to the same replicate were concatenated, MinION reads were de-multiplexed with cutadapt (Martin, 2011) and Illumina reads were clustered as described in the next section.

### Clustering of V4 hyper-variable region

Two different pipelines were used to cluster the V4 hyper-variable region: a distance-based method and an error-based method, depending on the sequencing nature of the sequences. The V4 region of sequences coming from Sanger and MinION sequencing was extracted with cutadapt (Martin, 2011) using the primer set used to amplify the V4 region (see ‘Amplification and sequencing’ section: TAR-EukF1 / TAR-EukR3) and allowing 10% of mismatch for Sanger sequences and 20% of MinION sequences. Sequence extracts were later clustered using swarm (Mahé et al., 2015) with 1, 2 and 3 differences. Reads coming from Illumina sequencing were clustered into ASVs using DADA2 and the pipeline described in Callahan et al. (2016). Final amplicons were compared against PR2 v4.14.0 (Guillou et al. 2013). Note DADA2 was not used for Sanger sequences or MinION output since the error-correction algorithm relies on number of reads.

### Consensus sequence building

Sanger sequences were aligned using MAFFT v7.395 (Katoh and Standley, 2013) with the G-INS-i algorithm (‘--globalpair’) and 1000 iterative refinement cycles. Final consensus sequences were produced with a custom script (see ‘alignmentConsensus.py’ in github.com/MiguelMSandin/IntraGenomic-variability) using a 70% consensus threshold and a 30% threshold for base detection. MinION sequences taxonomically assigned to Polycystines by BLAST were used to build consensus sequences according to Wurzbacher et al. (2019), using an OTU clustering cutoff of 30% due to the bad quality of the reads with the script *consension* (https://microbiology.se/software/consension/). Resulting consensus sequences from MinION were mapped against the raw fastq file using minimap2 (Li, 2018) and lastly polished with racon v1.4.22 (Vaser et al. 2021).

### Phylogenetic analysis

Consensus sequences coming from MinION and Sanger sequencing were aligned against a reference alignment using MAFFT v7.395 (Katoh and Standley, 2013) with a G-INS-i algorithm (‘--globalpair’) and 1000 iterative refinement cycles. The reference alignment is composed of reference sequences (extracted from Decelle et al., 2012; Biard et al., 2015; Sandin et al. 2019 and Sandin et al. 2021; Supplementary material Table S1) and consensus sequences obtained by Sanger sequencing in this study. Automatic alignment outputs were manually checked and corrected in AliView v1.26 (Larsson, 2014). Final alignments were trimmed automatically using trimal (Capella-Gutiérrez et al., 2009) with a 30% gap threshold. The final data set contains 525 taxa and 2434 positions. A phylogenetic analysis done by RAxML and GTR+Gamma and GTR+CAT evolutionary models with 1000 rapid bootstraps was used to infer the phylogenetic position of the consensus sequences. Final trees were visualized and edited with FigTree version 1.4.3 (Rambaut 2016).

### Entropy analysis

In order to compare different sequencing methods and discriminate errors from intra-genomic variability, we perform an entropy analysis. Sequences taxonomically assigned to Nassellaria and Spumellaria were aligned independently using MAFFT v7.395 (Katoh and Standley, 2013) with a G-INS-i algorithm and 1000 iterative refinement cycles for high accuracy. For every position of each alignment, Shannon entropy was calculated using a custom script (see ‘alignmentEntropy.py’ in github.com/MiguelMSandin/IntraGenomic-variability), without considering insertions or deletions (“-“) due to the incompleteness of Nassellaria sequences. For Sanger and MinION sequences, a reference alignment was build including only Sanger sequences as previously described, and then MinION sequences were aligned to the reference alignment. Later Sanger sequences were removed in order to obtain 4 different alignments: Nassellaria from Sanger, Nassellaria from MinION, Spumellaria from Sanger and Spumellaria from MinION. Further entropy values were calculated for the 4 alignments, after removing columns with only gaps. The entropy of the full V4 region was measured in an independent analysis to compare the three different sequencing technologies. The full V4 region was extracted as previously described (cutadapt) from Sanger and MinION sequencing results and sequences assigned to Nassellaria and Spumellaria from the PR2 v4.14.0 (after removing highly divergent sequence assigned to Orosphaeridae and Collodaria and duplicated sequences). Final datasets of Nassellaria and Spumellaria independently included sequences from 4 different origins: reference sequences from PR2 v4.14.0 and sequences obtained in this study by Sanger, MinION and Illumina sequencing.

## Results

### Quality of sequencing results

In total 834 reads (representing 737,817 bases) were successfully retrieved from Sanger sequencing for the 4 holobionts (from the total 864 sequences sequenced: 4 holobionts, 3 replicates per holobiont, 24 sequences per replicate and 3 reads (primers) per sequence; see Supplementary material Fig S1 for a schematic representation). These sequences had an average length of 884.67 (± 198.03) bp, with a median of 1005 bp and an average BLAST similarity identity of 98.13 (± 2.59) % with a reference sequence (Fig. 1, Sanger).

**Figure 1.**
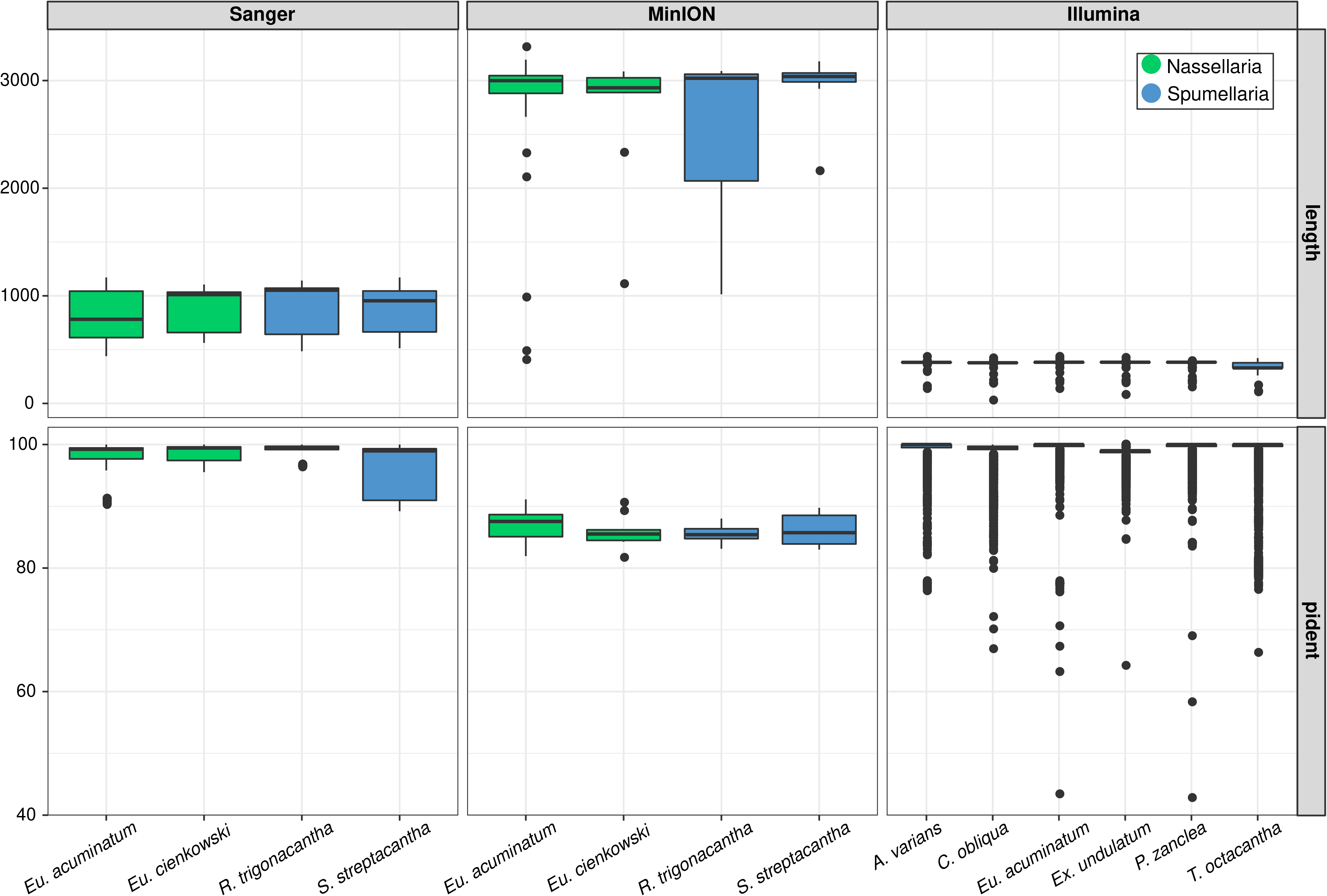
Boxplot summarizing sequencing results of Sanger, Oxford Nanopore Technologies (MinION) and Illumina sequencing; top row shows the sequence length (in base pairs; bp) and bottom row the percentage identity of the first match against a reference sequence per cell. Note that identity for Sanger and MinION was performed by BLAST and for Illumina by global alignment (vsearch) due to the shorter length of the reads.

MinION sequencing resulted in 864 total sequences (representing a total of 1,645,774 bases). The low throughput obtained during the last 28h of the flowcell live (20h → 48h) contrast with the high number of sequences obtained during the first 20h (0h → 20h) in an independent experiment, where there were a total of 225,573 sequences (representing 296,072,423 total bases; data being analyzed). Approximately, during the last 28h of the flowcell there were sequenced 0.38% of the total sequences (or 0.56% of the total bases) obtained by the same flowcell. These 864 sequences were compared against NCBI database by BLAST tool and 81 had no possible matching, 185 were assigned to bacteria, 593 to eukaryotes and 5 matching different domains (e.g. the same sequence was matching virus, uncultured prokaryote and uncultured organism). The remaining 783 sequences had an average sequence length of 2043.48 bp (± 1143.42) bp, with a median of 2746 bp and an average identity of 86.01 (± 2.91) % with a reference sequence. De-multiplexing MinION sequences resulted in 55 sequences among the different replicates. After de-multiplexing, the average sequence length increased to 2641.82 bp (± 681.80) bp and the median length to 2921 bp although the average identity remained at values of 86.48 (± 2.33) % similarity on average with a reference sequence (Fig. 1, MinION).

Regarding Illumina sequencing, a total of 1,019,196 sequences (representing 254,799,000 bases) were sequenced in both the R1 and the R2 files. On average 12,135 (± 6,470) reads were obtained for each replicate. These reads were merged resulting on 10,256 (± 6,278) amplicons on average per replicate and 66,913 total unique amplicons. They had an average length of 375.3 (± 15.89) bp, a median of 382 bp and an average identity of 99.28 (± 2.26) % with a reference sequence (Fig. 1, Illumina).

### Taxonomic assignation and diversity of reads

Sanger sequenced amplicons of the different parts of the ribosomal genes belonging to the same replicate were concatenated resulting in 287 sequences with an average length of 2570.7 (± 296.97) bp. This covers about the first ~1000 bp of the 18S rDNA gene (primer SA), the last ~600-800 bp of the 18S and the ITS1 rDNA gene (primer S69f) and the regions D1 and D2 from the 28S rDNA gene (primer D1R).

In *Eucyrtidium cienkowski* (Vil325) sequences were assigned to Nassellaria (56), Ciliates (11) and Tunicates (4); in *Eu. acuminatum* (Vil496) to Nassellaria (53), Chrysophyceae (14) and diatoms (5); in *Rhizosphaera trigonacantha* (Mge17-81) to Spumellaria only (72) and in *Spongosphaera streptacantha* (Mge17-82) the community was the most diverse with Spumellaria (63), fungi (5), ciliates (Apicomplexa: 2) and dinoflagellates (1) (Fig. 2, Sanger). From the de-multiplexed sequences extracted from MinION sequencing, in *E. cienkowski* was found only Nassellaria (9); in *E. acuminatum* was found Nassellaria (16), diatoms (6), chrysophyceae (4) and fungi (1); in *R. trigonacantha*, only Spumellaria (8), as in Sanger sequencing, and in *S. streptacantha* a similar composition as in Sanger, Spumellaria (7), dinoflagellates (3), alveolates (1) and fungi (1) (Fig. 2, MinION).

**Figure 2.**
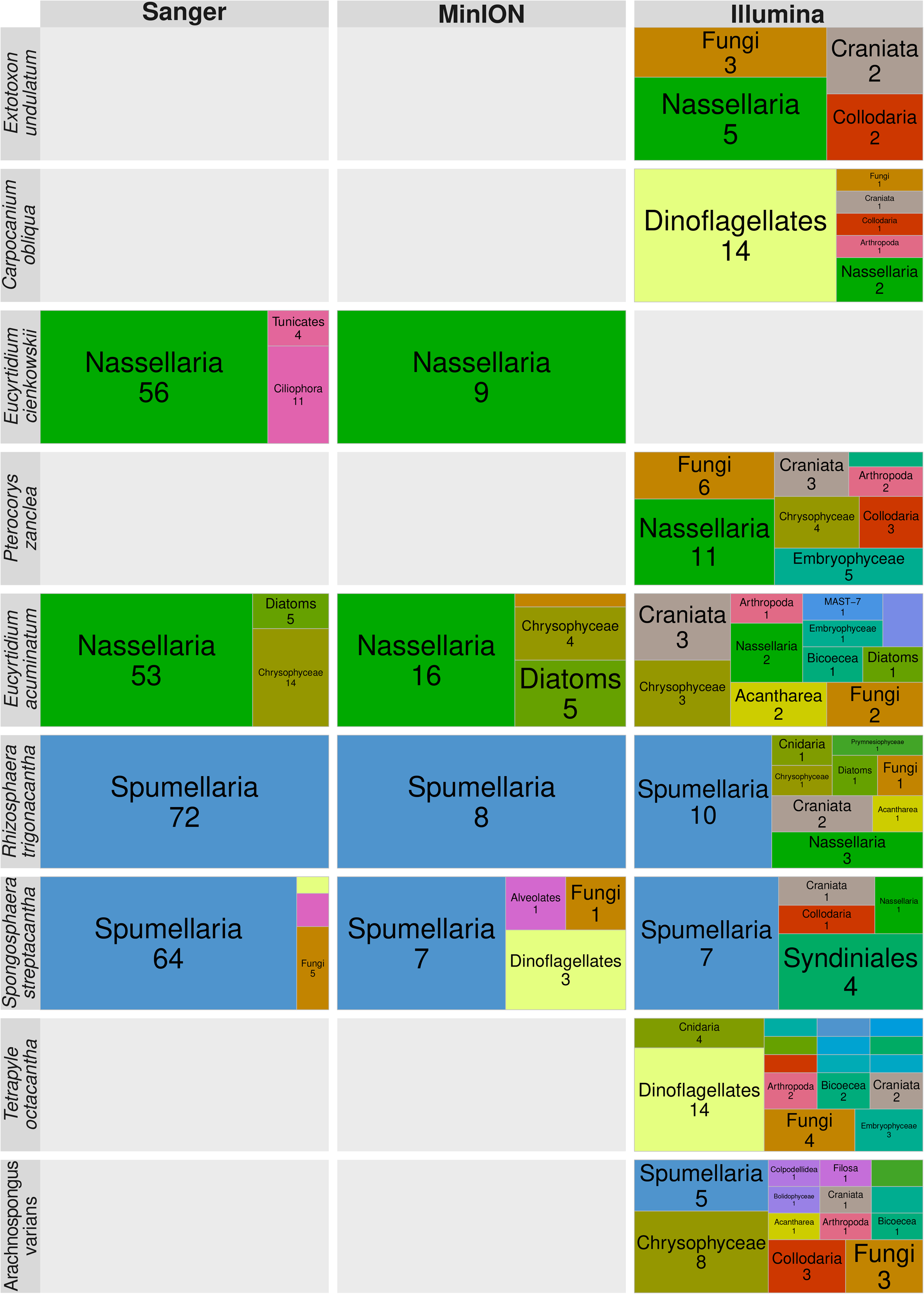
Taxonomic affiliation of sequences/amplicons obtained by Sanger (after concatenation of the primers), Oxford Nanopore Technologies (MinION) and Illumina (reads processed by dada2) sequencing for each cell. The area represents the proportion of total number of unique sequences/amplicons affiliated to the specific taxonomic entity (tree map). Numbers below the taxonomic group represent the number of unique sequences/amplicons.

Amplicons of Illumina were clustered using an error-based method (dada2), in total 153 ASVs (with a total of 570,041 reads) were found. These ASVs had an average length of 379.2 (± 8.99) bp, a median length of 378 bp and an average similarity identity of 99.70 (± 5.97) %. The most diverse groups are dinoflagellates (28 ASVs), Spumellaria (23 ASVs), Nassellaria (19 ASVs) and Fungi (14 ASVs mostly within Agaricomycotina and Ustilaginomycotina) (Fig. 2, MinION). Despite the big diversity within each holobiont, the number of reads is highly dominated by 1, 2 or 3 ASVs belonging to the host or the symbiotic algae (Supplementary material Fig S3). Due to the great taxonomic diversity found within holobionts by Illumina sequencing, only ASVs present in at least 3 samples (triplicates in the PCR), with a total abundance equal or higher than the median value of the abundance (that is 102 reads) and assigned until at least the taxonomic rank “Order” were considered. We have chosen stringent thresholds in an attempt to remove all artefacts and/or contaminations, understanding that part of the targeted diversity within the holobionts might also be removed. After processing, the number of ASVs decreased to 36 ASVs but the total reads did not change drastically with 544,127 reads, representing up to 95.45% of the total reads. Main taxonomic affiliations were similar to the one described before filtering the ASVs. Furthermore, the relative proportion of unexpected ASVs took more importance; such as an ASV affiliated to Collodaria present in 4 holobionts, an ASV affiliated to Acantharea present in 3 holobionts and 1/2 ASVs affiliated to Craniata present in every sample (Supplementary material Fig S4). When exploring in deeper detail the ASVs affiliated to Polycystines and their abundance and distribution among samples after applying the stringent thresholds, it is possible to find up to 3 different and highly abundant ASVs within the same cell (e.g. *Spongosphaera streptacantha* or *Arachnospngus varians*; Fig. 3). Some of the most abundant ASVs were also found in a fourth and fifth sample; as for “ASV-2”, “ASV-5” and “ASV-7”, ASVs from the Nassellaria hosts *Pterocorys zanclea*, *Extotoxon undulatum* and *Eucyrtidium acuminatum* respectively. “ASV-2” was present in the three replicates of *E. acuminatum* and in one replicate of *Carpocanium obliqua* and *Rhizosphaera trigonacantha*, a Spumellaria host, although at very low relative abundances. In *E. undulatum* there were two ASVs, of which one (“ASV-5”) was present in the three replicates of the holobiont and in a fourth and fifth replicate from *Rhizosphaera trigonacantha* and *Spongosphaera Streptacantha*, the second ASV (“ASV-47”) was only present in one replicate. This ASV, is also found in *C. obliqua*, *T. octacantha* and *A. varians*, it never appears within the same three samples from an independent holobiont and its taxonomic affiliation is to Collodaria, with a 100% similarity to a reference sequence and 100 BS support for the taxonomic assignation.

**Figure 3.**
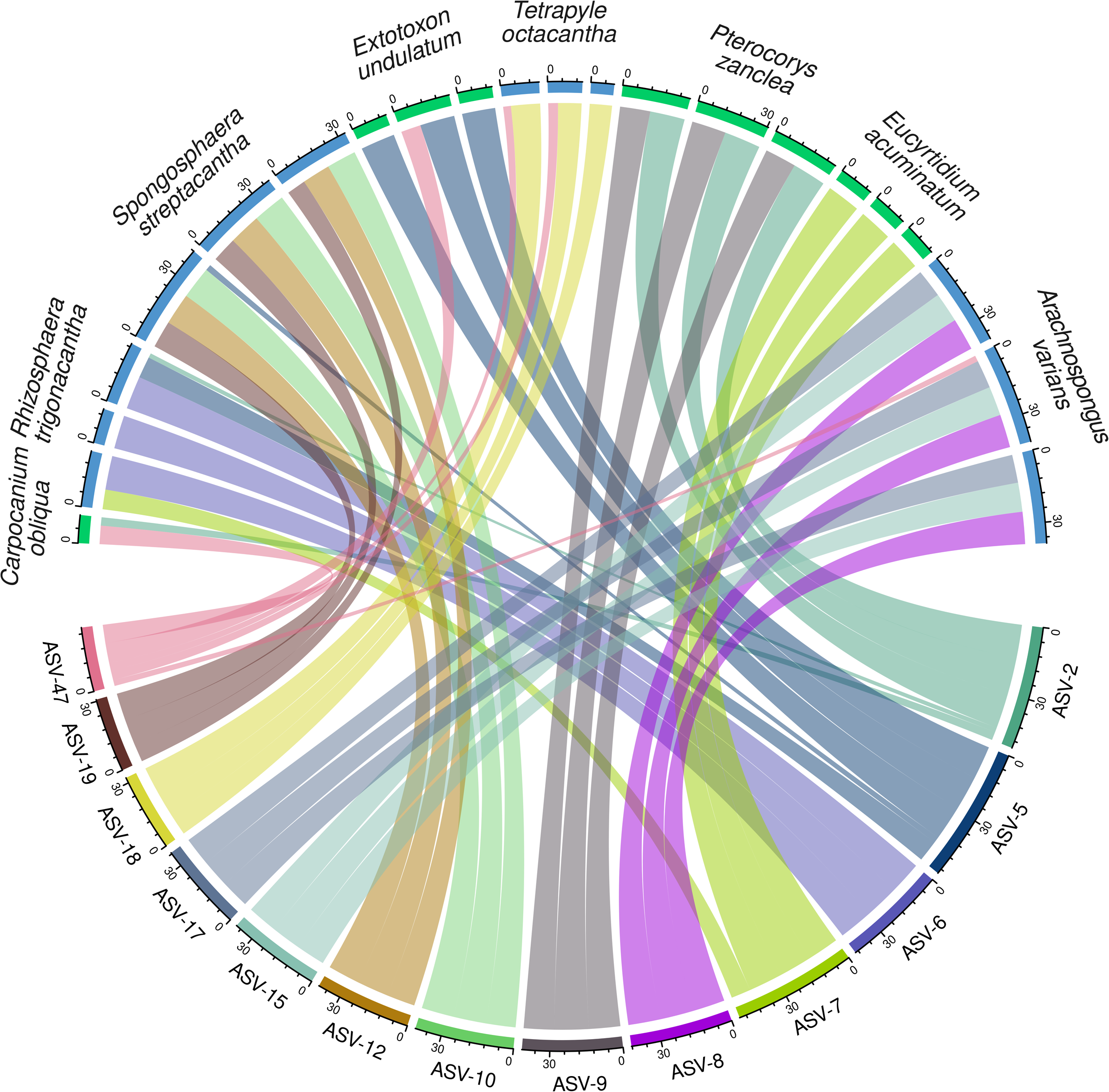
Circular plot representing the log transformed abundance of polycystines amplicons (lower half of the circles) clustered with dada2 and sample affiliation (upper half of the circles). Only amplicons affiliated to polycystines, present in 3 or more samples and with a total abundance equal or higher than 102 reads (median) were considered.

### Intra-genomic variability

The hyper-variable region V4 of the 18S rDNA gene was extracted from the sequences affiliated to Radiolaria from both Sanger and MinION sequencing, in order to compare with Illumina sequencing and assess if it is detectable the same genetic diversity within the holobiont. In total 52, 56, 59 and 68 sequences from Sanger sequencing were extracted belonging to *Eucyrtidium acuminatum*, *E. cienkowski*, *Spongosphaera streptacantha* and *Rhizosphaera trigonacantha* respectively. The same unique sequence was found to be the most abundant in both *E*. *acuminatum* and *E. cienkowski* (54 and 47 reads respectively) being exactly identical despite of being morphologically identified in different species, and up to 5 other unique sequences were found with a very low abundance (2, 2, 1, 1 and 1 reads, green dots on Fig. 4 under unique reads). These sequences were very similar, showing similarities within them close to 100% with a maximum of one base of difference (boxplots on Fig. 4 under unique reads). Similar patterns are seen in the Spumellaria *R. trigonacantha*, with one sequence found 67 times and only one sequence different with one read. In contrast, *S. streptacantha* shows up to 3 different reads relatively abundant (16, 13 and 13 reads) and 8 other sequences with a lower abundance, having a similarity among them of 96.1%. It is important to note, that these three sequences were sequenced in the three different replicates (PCR reactions). When these sequences were clustered with swarm and 1 difference of threshold, *Eu*. *acuminatum* and *Eu. cienkowski* share the same amplicon. In *R. trigonacantha* the single sequence was grouped in the unique amplicon found for this holobiont, and up to 3 different amplicons were found in *S. streptacantha* (still with a similarity among these amplicons of ~96%). Increasing the difference threshold to 2 differences the amplicons in *S. streptacantha* remain comparable to 1 difference. When using a clustering of 3 differences, the number of amplicons in *S. streptacantha* decreases to 2, with low changes in the similarities among them (96.4%, representing 14 different aligned positions in 385 an 382 bp sequences long). Yet one amplicon largely dominates the other (51 reads against 8 reads). Same protocol was implemented in MinION sequences in order to extract the V4 hyper-variable region of the 18S rDNA (data not shown). In total 4 sequences were extracted from *R. trigonacantha*, 2 from *S. streptacantha*, 6 from *Eu. cienkowski* and 20 from *Eu*. *acuminatum*. It was not possible to cluster the sequences with swarm due to their large dissimilarities. Up to a difference threshold of 10, there were still no clusters, keeping an average intra-genomic similarity of 86.92% (± 3.56%).

**Figure 4.**
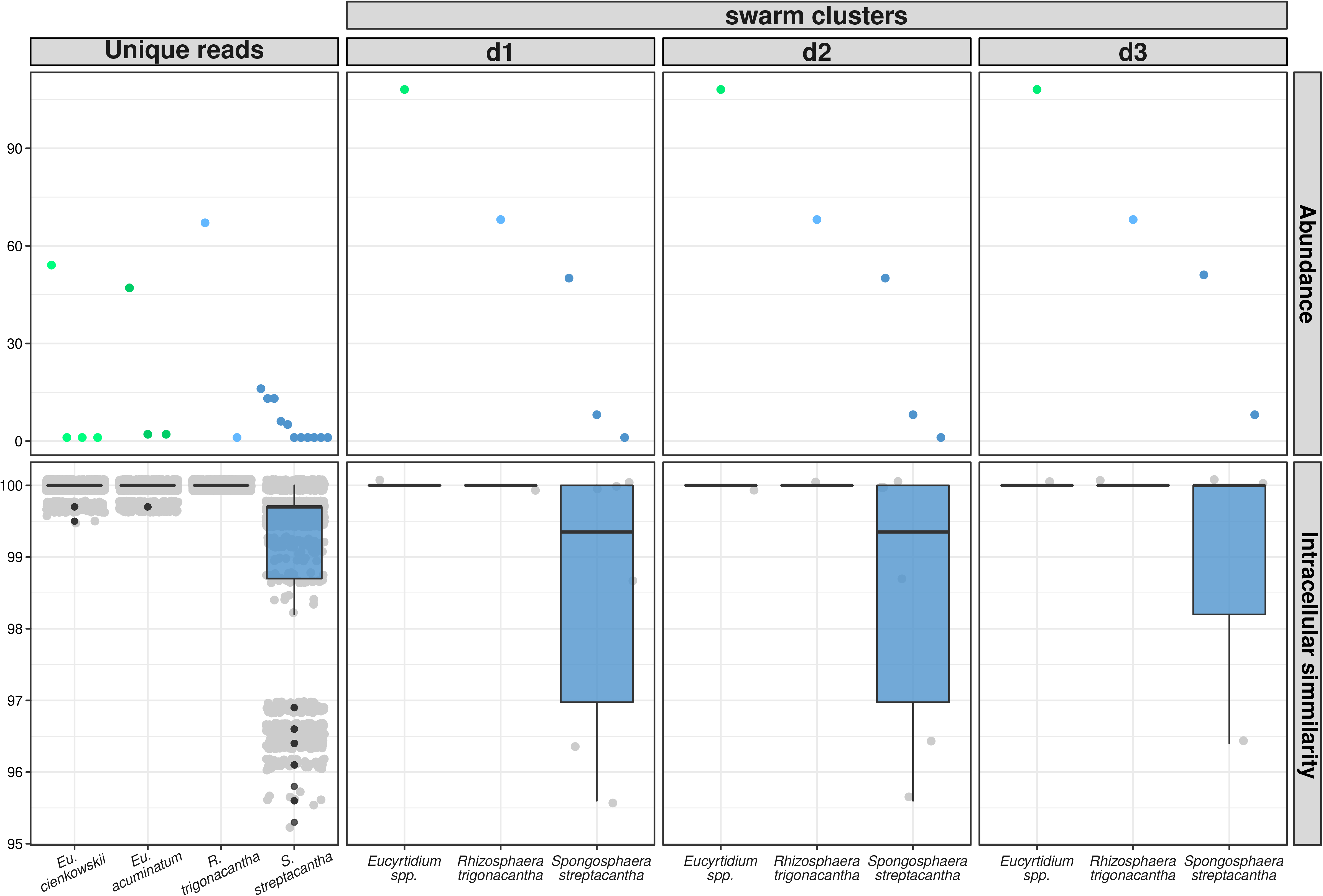
Abundance and intra-genomic similarities of the V4 region extracted from Sanger sequencing results of the total unique reads and clustered with swarm with 1 (d1), 2 (d2) and 3 (d3) differences.

In order to compare the intra-genomic variability between different sequencing methods and discriminate between sequencing errors, we calculated the entropy of the alignment from sequences obtained in this study at every position. Since it is expected that errors are random, a single error in the alignment would have low evenness and appear with low entropy (yet not 0). On the other side, intra-genomic variability is expected to be sequenced in several replicates increasing the evenness and with it the entropy, and also with a tendency to appear towards hyper-variable regions. In general, concatenated sequences from Sanger sequencing belonging to Nassellaria showed low Shannon entropy values among them (Supplementary material Fig S5). Towards the end of the primers the entropy slightly increased, meaning that most probably are suspected to be errors. Especially at the end of the region D2 of the 28S rDNA gene the entropy reaches its highest values along a region of ~100 bp length, probably meaning variability among the different copies of the rDNA. Regarding Spumellaria, *Rhizosphaera trigonacantha* has a similar trend than Nassellaria (Supplementary material Fig S5). In contrast, *Spongosphaera streptacantha* shows regions of high entropic values at around the position 750 (V4 region from the 18S rDNA gene), along the beginning of the 28S RDNA and especially over a region of ~250 bp on the ITS1, showing a most probable big intra-genomic variability of the rDNA. In general MinION sequences aligned against the reference sequence with many gaps (Supplementary material Fig S5). Alignments of sequences coming from MinION have in general a greater number of positions than alignments of sequences coming from Sanger sequencing (~922.25 bp more in average); greater than the length of the 5.8S and ITS2 regions of the rDNA that MinION sequences are covering in comparison to Sanger sequences (18S+ITS1 and D1 and D2 regions of the 28S rDNA). These sequences had high entropy values all along the rDNA alignment. With the exception of the 18S rDNA gene of Nassellaria, where there is a constant entropy trend, there are variations of the entropy depending on the region, but these variations do not match exactly those of the Sanger sequences.

The V4 hyper-variable region of the 18S rDNA gene used in Fig. 4 along with that extracted from MinION sequencing and Illumina were pooled together in one alignment for the V4 region comparison. In total, 108 sequences for Nassellaria and 127 sequences for Spumellaria were extracted from Sanger sequencing, 19 Nassellaria sequences and 5 Spumellaria sequences were extracted from MinION sequencing, including those amplicons extracted from Illumina taxonomically assigned to Nassellaria (19) and Spumellaria (23) and completed with reference sequences from pr2 v4.14.0 (83 for Nassellaria and 671 for Spumellaria) trying to gather most of their known diversity. Final aligned dataset had a length of 478 bp for Nassellaria and 461 bp for Spumellaria (Fig. 5). Reference sequences showed a hotspot of higher Shannon diversity within the hyper-variable region in both Nassellaria and Spumellaria spanning from the position ~80 until ~210 with maximum values around the position ~110-150. Nassellaria reference sequences showed two other hotspots regions peaking at around the positions ~330 and ~420, yet average entropy values are half of the first hotspot region. Regarding Spumellaria, these two last hotspot regions are less marked than for Nassellaria due to overall higher entropy values. Sanger sequences maintain a near-0 entropy values, with the exception of a region between ~100-200 bp in Spumellaria showing higher entropy values (corresponding to *Spongosphaera streptacantha*). Illumina ASVs follow same patterns as the reference alignment, with the highest entropy values at around the position ~150. Despite the poor trends found for the full length of the rDNA for MinION sequences (Supplementary material Fig S5), when focusing on the V4 region, shows similar patterns that those found in the reference alignment and Illumina, although these peaks are smoother due to the relative higher entropy values all along the alignment.

**Figure 5.**
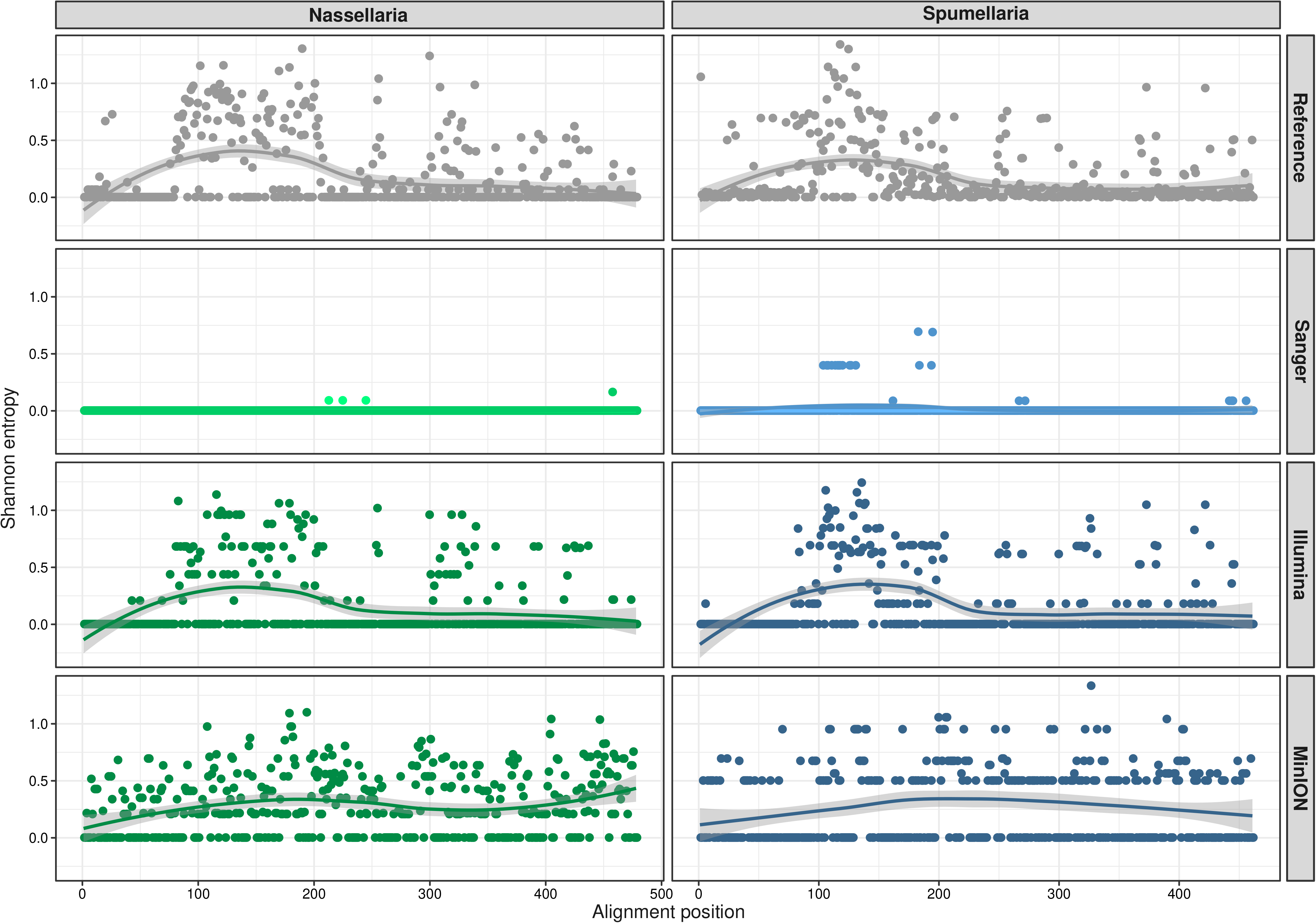
Shannon entropy analysis for every position (on *x* axis) of the V4 hyper-variable region of the 18S rDNA gene for reference sequences of Nassellaria and Spumellaria (extracted from PR2 v4.14.0) and for Sanger, Illumina and Oxford Nanopore Technologies (MinION) sequencing results aligned all together within the different taxonomic groups. For further results on the near-full rDNA alignment entropy obtained by Sanger and Oxford Nanopore Technologies (MinION) sequencing see supplementary figure S5.

We performed a last attempt to test whether the high-error rate from MinION is affecting further downstream analysis or is in the contrary possible to be addressed. Sequences coming from Sanger and MinION were clustered into consensus sequences and were aligned in a phylogenetic tree along reference sequences (Supplementary material Table S1). Despite the high error rate of MinION sequences, the random distribution of the errors and the long reads bring accurate phylogenetic information when processed into consensus sequences (Fig. 6). Consensus sequences from MinION are phylogenetically sister to the consensus sequences coming from Sanger showing high phylogenetic support and short branches. Consensus sequences from Sanger sequencing have a similarity >99% against reference sequences except for Eucyrtidium cienkowski (97.32%). Consensus sequences from MinION have a maximum similarity identity against reference sequences of 99.5%, 98.9% and 99.2% for *Eucyrtidium* sp., *Rhizosphaera trigonacantha* and *S. streptacantha* respectively (data not shown).

**Figure 6.**
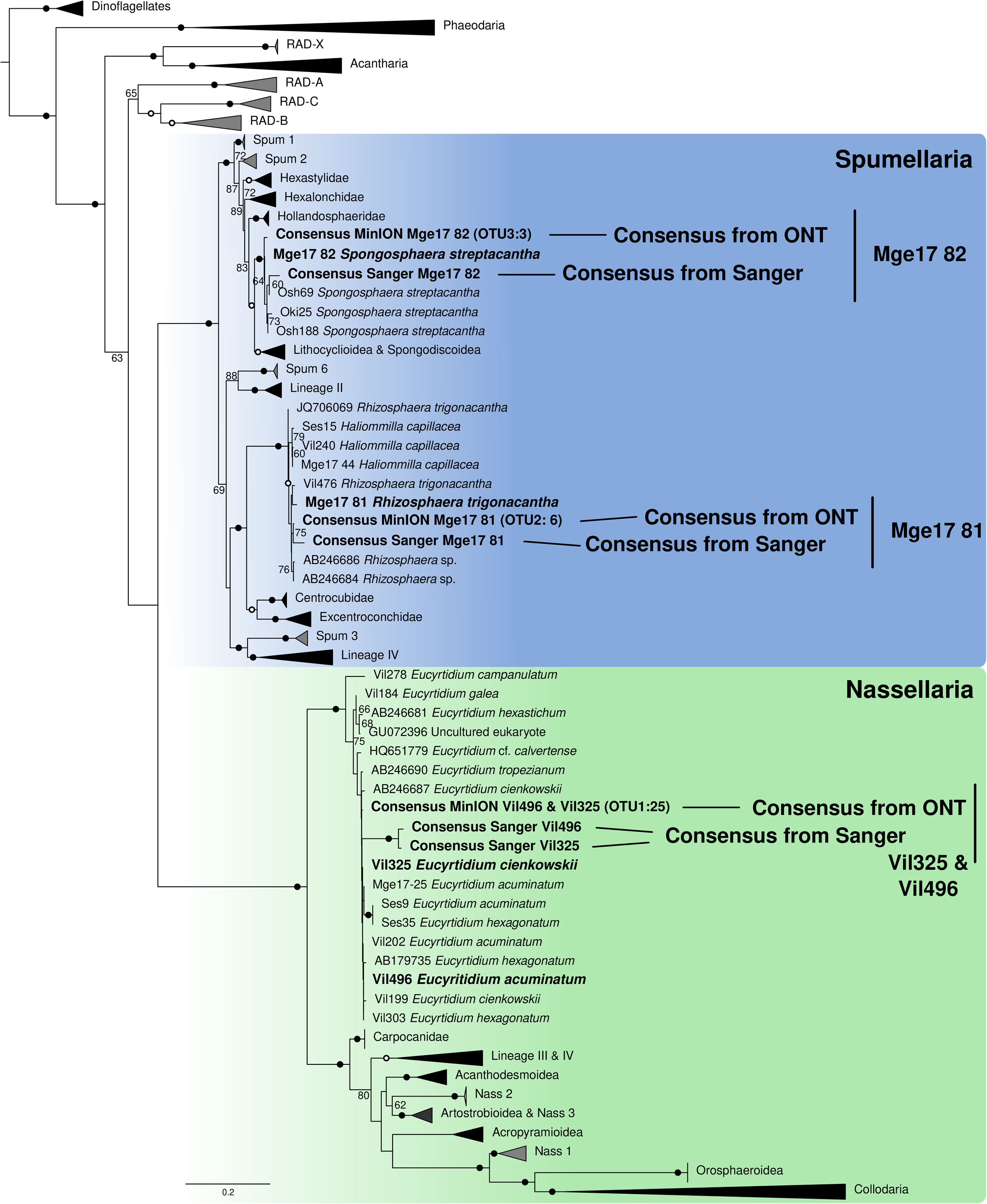
Molecular phylogeny of consensus Oxford Nanopore Technologies (MinION) sequences, consensus of Sanger results and reference sequences inferred from the concatenated complete 18S and partial 28S (D1-D2 regions) rDNA genes. In bold are shown specimens used in this study (consensus) and reference sequences previously obtained for the same specimens. Numbers after the OTU name of sequences obtained by MinION represent from how many raw sequences the consensus was built. The tree was obtained by using a phylogenetic Maximum likelihood method implemented in RAxML using the GTR + CAT model of sequence evolution and 1000 rapid bootstraps (BS, shown at the nodes). Bootstrap values below 60 are not shown. Black circles indicate BS ≥ 99%. Hollow circles indicate BS ≥ 90%.

## Discussion

Our results showed that intra-genomic rDNA gene variability of Nassellaria and Spumellaria is generally limited, showing in some cases the same V4 rDNA hyper-variable region between different morpho-species (yet belonging to the same genus; *E. cienkowskii* and *E. acuminatum*), as previously found in tintinnids ciliates (Bachy et al., 2013). Although in some species intra-genomic rDNA gene variability can be very important, as it is the case of *Spongosphaera streptacantha* in which we found two distinct intra-genomic V4 rDNA regions (~97% similarity, representing 14 different nucleotide positions). Such taxonomic differences among closely related groups have also been found in Oligotrich and Peritrich Ciliates (Gong et al., 2013). Most of this intra-genomic variability is however overlooked due to the presence of a highly repeated copy that predominates over the low abundant copies, as previously found in Nassellaria or among other orders of Radiolaria such as Acantharia (Decelle et al., 2014). However, in Acantharia the intra-genomic variability could also become important, finding up to 3 different OTUs (V9: clustered at 97% and present in 2 replicates; Decelle et al., 2014). Similar studies have shown a relationship between the intra-genomic variability and the number of macronuclei (Zhao et al., 2019) and between the rDNA copy number in ciliates (Gong and Marchetti, 2019) and in alveolates (Medinger et al., 2010). That could explain the taxonomic differences in the intra-genomic variability of Radiolaria, since both Nassellaria and Spumellaria have only one nucleus that tend to be small whereas Acantharia and Collodaria have several nuclei (Suzuki et al., 2009). In the former case, they tend to show low intra-genomic variability, whereas in the later case can be relatively high (Decelle et al., 2014).

Despite the low intra-genomic variability found within the hosts, Illumina sequencing has identified *a priori* a very diverse host gene variability; finding in some hosts up to 10-11 different ASVs (Fig. 2). And when comparing the holobiont community obtained by the three different sequencing methods, Illumina shows a notably diverse holobiont community. The unexpected presence of taxonomic groups such as Acantharia, Collodaria or Craniata within holobionts constituted by Nassellaria and Spumellaria as hosts, questions the reliability of the so-called “rare” ASVs in environmental studies. Part of this rare biosphere has been suggested as artefacts inflating diversity estimates (Kunin et al., 2010; Bachy et al., 2013). Other ASVs present in a fourth or fifth sample (i.e.: “ASV-2”, “ASV-5” and “ASV-7” in Fig. 3) also question technical issues such as cross-contamination during sequencing (Kircher et al., 2012) or tag-jumping during library preparation (Schnell et al., 2015). Other ASVs have had full similarities against a reference sequence and have passed the stringent abundance filters we have applied (e.g. Craniata in fig. 2 or “ASV-47” affiliated to Collodaria in fig. 3), suggesting the presence of environmental DNA (eDNA) since all holobionts were collected in the north-western Mediterranean Sea, yet at varying depths, localities and dates. Numbers of reads have been found to be correlated with the number of nuclei and the cell size (Biard et al., 2017; Pitsch et al., 2019). Our results indicate that eDNA contamination is showing a bias towards organisms with a higher copy number, such as Collodaria (Biard et al., 2017) or the metazoans Craniata. A problem that might be accentuated when the targeted DNA is relatively low abundant (*i.e.* that of Nassellaria) and have to “compete” for the available space in the flow-cell during sequencing, resulting in large differences of relative abundances, as seen in *Carpocanium obliqua* and *Tetrapyle octacantha* (Fig. 2 and supplementary figure S4). Probably, DNA from Acantharia and Collodaria are more prone to be successfully amplified regarding that of Nassellaria and Spumellaria, as seen in differences of DNA amplification success among distinct Foraminifera taxa (Weiner et al., 2016). In this study we have used general eukaryotic primers that are equally binding these radiolarian groups. Therefore, further analysis must assess the effect on differences in shell or cell architecture regarding DNA extraction and amplification.

These results highlight the need for a careful interpretation of environmental metabarcoding surveys due to the taxonomic bias found among different taxa (e.g. Weiner et al., 2016; Gong and Marchetti, 2019). In some cases, illumina sequencing from eDNA tends to lead to an under-representation of the environmental diversity (e.g.: *Carpocanium obliqua* and *Tetrapyle octacantha* in Fig. 2) and to an over-estimation for some other taxa (e.g.: Collodaria and Craniata). Most of these biases and technical problems cannot be identified and sorted out in environmental metabarcoding surveys, most frequently due to the lack of replicates, leading to poor estimations of the environmental molecular diversity. For example, the presence of “ASV-2”, “ASV-5” or “ASV-7” in a fourth and fifth sample would have been considered as low abundant ASVs in those specific samples in global environmental metabarcoding surveys lacking replicates. This emphasizes the need for considering replication in metabarcoding surveys to ensure an accurate estimation of the environmental microbial diversity (Prosser, 2010), being technical replicates more important than biological replicates. Besides, understanding other aspects may help the correct interpretation of metabarcoding surveys, especially important when working with specific taxa, such as the estimation of the rDNA copy number (Biard et al., 2017; Gong and Marchetti, 2019), the analysis of differences in relative abundance (Morton et al., 2019) or even exploring previous steps such as DNA extraction bias, amplification primers or intracellular architecture. However, a big part of the overestimated diversity might also be due to sequencing errors, as previously proposed (Bachy et al., 2013; Decelle et al., 2014), since the 50% less abundant ASVs accounted for less than 5% of the total reads meaning a high presence of singletons and low abundant clusters.

Illumina sequencing showed to be the method with the highest sequencing depth. After clustering into ASVs, the error rate showed to be the lower from the three methods and randomly distributed. Besides, most of the suspected artefacts produced during Illumina sequencing were removed applying stringent filters. Sanger sequencing also showed a limited error rate, yet singletons were found in the V4 rDNA region of every sample sequenced, and were later corrected when clustering (Fig. 4). However, towards the end of the primers the error rate increased substantially. For this reason, we suggest that filtering of Illumina ASVs should be done more importantly based in abundance than identity thresholds, especially for the V9 region that is located near the end of general and specific primers. On the other side, MinION is the technique with the highest error rate, finding alignment similarities of the raw reads around 94-97% in bacteria (Tyler et al., 2018) better than our results with an average of 86% for the best scoring sequences. Although, recent studies have generated consensus sequences decreasing considerably the error rate and reaching results comparable or better than those obtained by Sanger sequencing (Pomerantz et al., 2018; Wurzbacher et al., 2019), as seen in our study (>98.9% similarity identity to reference sequences). Despite the high error rate of raw reads from MinION found in our analysis, consensus sequences showed accurate phylogenetic signal with high bootstrap supports. Therefore it turns out that the length of the reads over-competes the randomly generated sequencing errors and brings accurate phylogenetic information. In our study we have used the last 28h of the MinION sequencing device, largely affecting the life and quality of the nanopores from the MinION flowcell in detecting the ion signal. It is therefore recommended to stop the flowcell according to dessired acquisition of reads and not time sequenced when using different projects in the same flowcell; as seen in Srivathsan et al. (2021) where they have successfully re-used a flow cell up to four times. Yet, when looking at the holobiont community sequenced in our study, the results obtained with MinION are comparable to those obtained by Sanger sequencing (Fig. 2).

ONT have the advantage of directly sequence the DNA strand with a theoretical no need of PCR amplification, and therefore removing biases associated in these steps that most likely produced part of the artificial environmental diversity found in Illumina. This also brings the possibility to work with absolute reads, and not relative values providing a more quantitative picture of the genetic community. Although rDNA copy number will still be an issue when comparing genetic and morphological diversity. ONT’s accuracy is improving since it was firstly released (Rang et al., 2018), and new methods for correcting and analysing ONT results are coming along (e.g.: demultiplexing: Wick et al., 2018; base-calling: Wick et al., 2019, Xu et al. 2020; consensus sequence building: Pomerantz et al., 2018; Wurzbacher et al., 2019; and polishing: Vaser et al. 2021). Therefore, MinION could generate fruitful results in the near future for metabarcoding surveys by taking advantage of the extensive work of the 18S reference sequences already done and the high variability and taxonomic resolution of the ITS and the 28S (as seen in environmental studies performed with PacBio, Jamy et al. 2019). The high cost-effectiveness of ONT (Cui et al. 2020) and the portability and sequencing depth and length of MinION device show fruitful perspectives for its implementation of environmental phylogenetic studies. Along with the high accuracy of circular consensus sequencing of PacBio (Wenger et al., 2019) for the high throughput establishment of reference sequences, environmental phylogenetic studies could move towards the full rDNA surveys.

## Acknowledgments

This work was supported by the IMPEKAB ANR 15-CE02-0011 grant and the Brittany Region ARED C16 1520A01. We would like to thank the MOOSE cruise and program for the opportunity of sampling and the facilities given on board, as well as John Dolan for hosting us multiple times at the Laboratoire d’Océanographie of Villefranche-sur-Mer. We are greatly thankful to Charles Bachy and Michal Karlicki for fruitful discussions on the intra-genomic variability and the analysis of Oxford Nanopore Technologies sequencing results, Nicolas Henry for help on the analysis and interpretation of Illumina sequencing results and Anna Karnkowska for constructive comments on earlier versions of this manuscript.

## Originality-Significance Statement

This work applies and asses the validity of a novel method for the exploration of the molecular environmental diversity as is the long-read sequencing from Oxford Nanopore Technologies, using the highly portable and cost-effective device MinION. In addition, our results provide a basis for the interpretation of already established short-read sequencing approaches for an ubiquitous and abundant marine protists, as is the Radiolaria.

## Supplementary Data

**Table S1**. List of publicly available sequences used in Fig. 6.

**Figure S1.** Schematic representation of the study design.

**Figure S2**. Light microscopy images of specimens used in this study. On the right of each specimen it is indicated whether it was used (tick mark) or not (cross mark) for Sanger + Oxford Nanopore Technologies (MinION) sequencing and/or Illumina, and in grey, a brief description on location and time of isolation. Scale bar (when available; black/white bar) represents 50 μm.

**Figure S3**. Tree map of the total number of reads (abundance) of every ASV affiliated to a taxonomic group for each cell obtained by Illumina sequencing. ASVs were processed by dada2.

**Figure S4**. Tree map of every ASV affiliated to a taxonomic group (left column) and their total number of reads (abundance; right column) for each cell obtained by Illumina sequencing. ASVs were processed by dada2. Only ASVs present in 3 or more samples and with a total abundance equal or higher than 102 reads (median) were considered.

**Figure S5**. Shannon entropy analysis for every position (on *x* axis) of Sanger and Oxford Nanopore Technologies (MinION) sequencing results aligned independently for Nassellaria (green) and Spumellaria (blue). Lines represent the tendency of the entropy for each alignment. Black arrows in Sanger boxes represent direction and approximate position of the primers used for Sanger sequencing. Insertions (“-“) were not considered due to the differences in length of the sequences and the high amount of insertions produced by the raw sequences from MinION.

