## Supplementary material for "Intra-genomic rDNA gene variability of Nassellaria and Spumellaria (Rhizaria, Radiolaria) assessed by Sanger, MinION and Illumina sequencing": media-2.pdf

### Nassellaria

| Specimen | Sanger+ MinION | Illumina |
|---|---|---|
| *Eucyrtidium cienkowski*<br>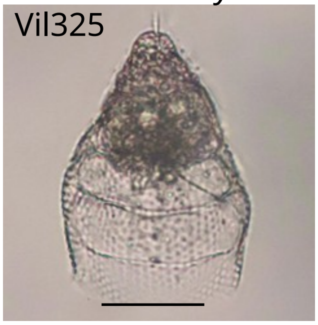<br>Villefranche-sur-Mer<br>43.681, 7.328<br>Surface<br>17/11/2012 | ✓ | ✗ |
| *Eucyrtidium acuminatum*<br>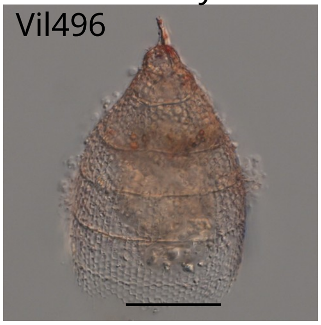<br>Villefranche-sur-Mer<br>43.681, 7.319<br>0-30m deep<br>23/10/2016 | ✓ | ✓ |
| *Extotoxon undulatum*<br>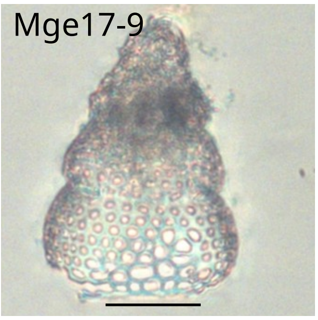<br>NW Mediterranean Sea<br>43.032, 5.199<br>0-500m deep<br>13/09/2017 | ✗ | ✓ |
| *Carpocanium obliqua*<br>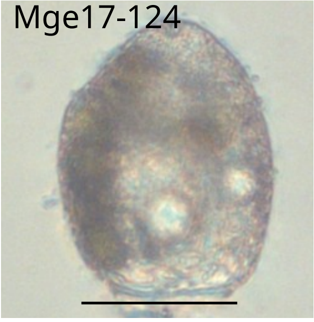<br>NW Mediterranean Sea<br>40.301, 6.283<br>0-500m deep<br>22/09/2017 | ✗ | ✓ |
| Pterocorys zanclea<br>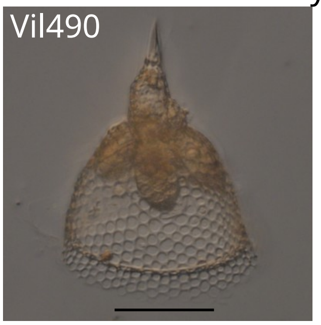<br>Villefranche-sur-Mer<br>43.681, 7.319<br>0-30m deep<br>22/10/2016 | ✗ | ✓ |

### Spumellaria

| Specimen | Sanger+ MinION | Illumina |
|---|---|---|
| *Rhizosphaera trigonacantha*<br>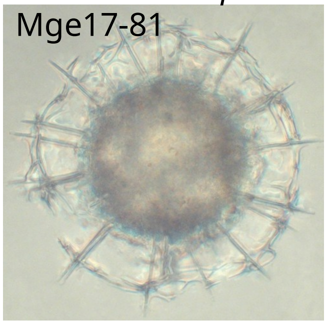<br>NW Mediterranean Sea<br>41.416, 6.45<br>0-500m deep<br>20/09/2017 | ✓ | ✓ |
| *Spongosphaera streptacantha*<br>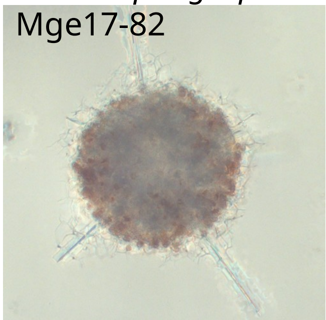<br>NW Mediterranean Sea<br>41.416, 6.45<br>0-500m deep<br>20/09/2017 | ✓ | ✓ |
| *Tetrapyle octacantha*<br>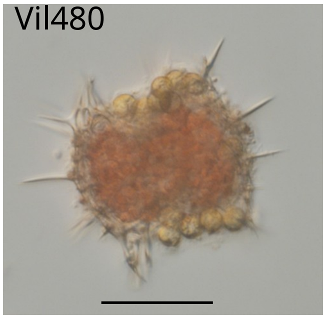<br>Villefranche-sur-Mer<br>43.681, 7.32<br>Surface<br>21/10/2016 | ✗ | ✓ |
| *Arachnospongus varians*<br>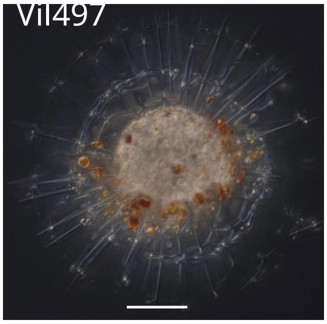<br>Villefranche-sur-Mer<br>43.681, 7.319<br>0-30m deep<br>23/10/2016 | ✗ | ✓ |
