## Supplementary figures and images for "Intra-genomic rDNA gene variability of Nassellaria and Spumellaria (Rhizaria, Radiolaria) assessed by Sanger, MinION and Illumina sequencing"

### media-1.pdf

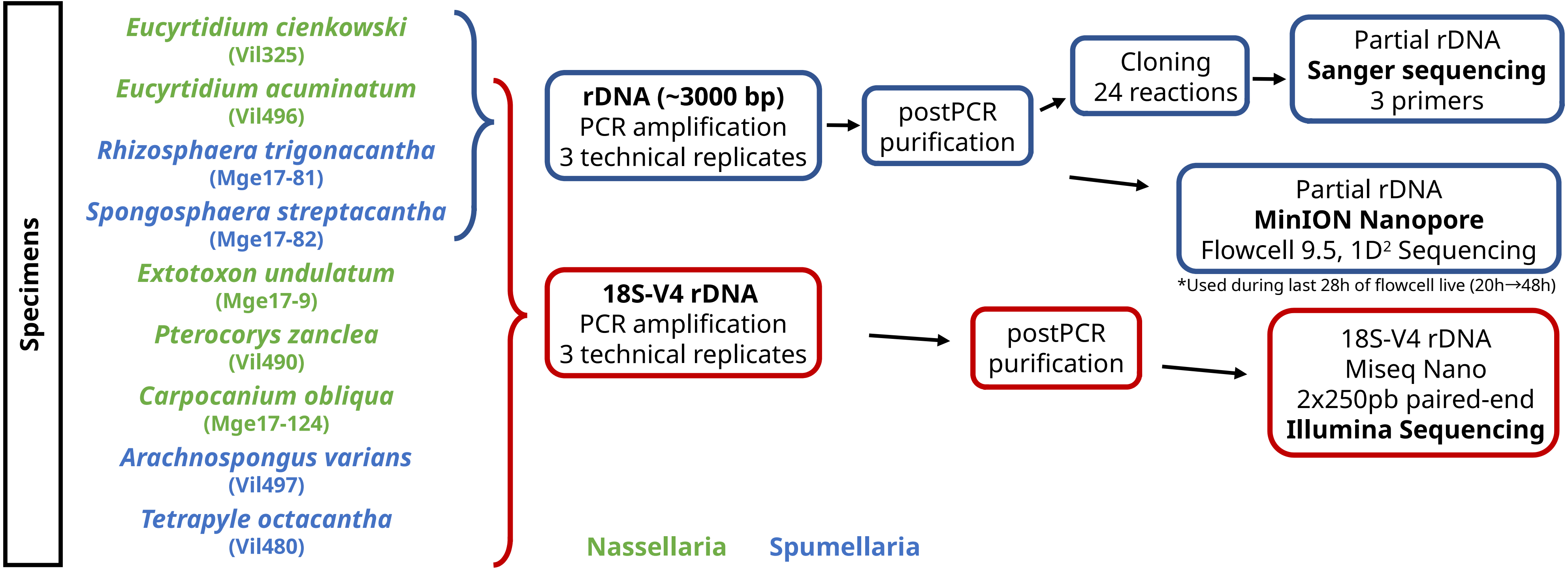

### media-3.pdf

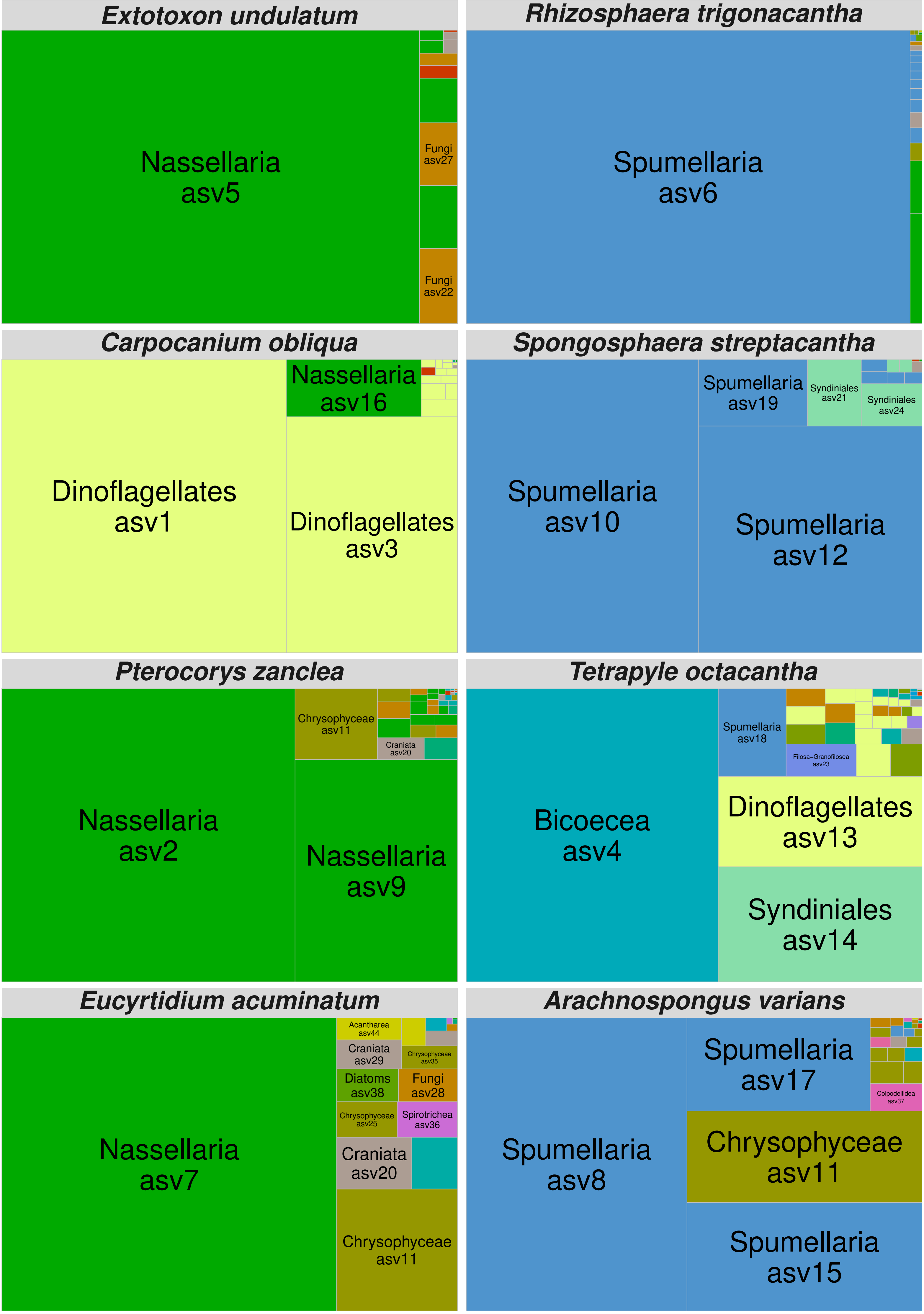

### media-4.pdf

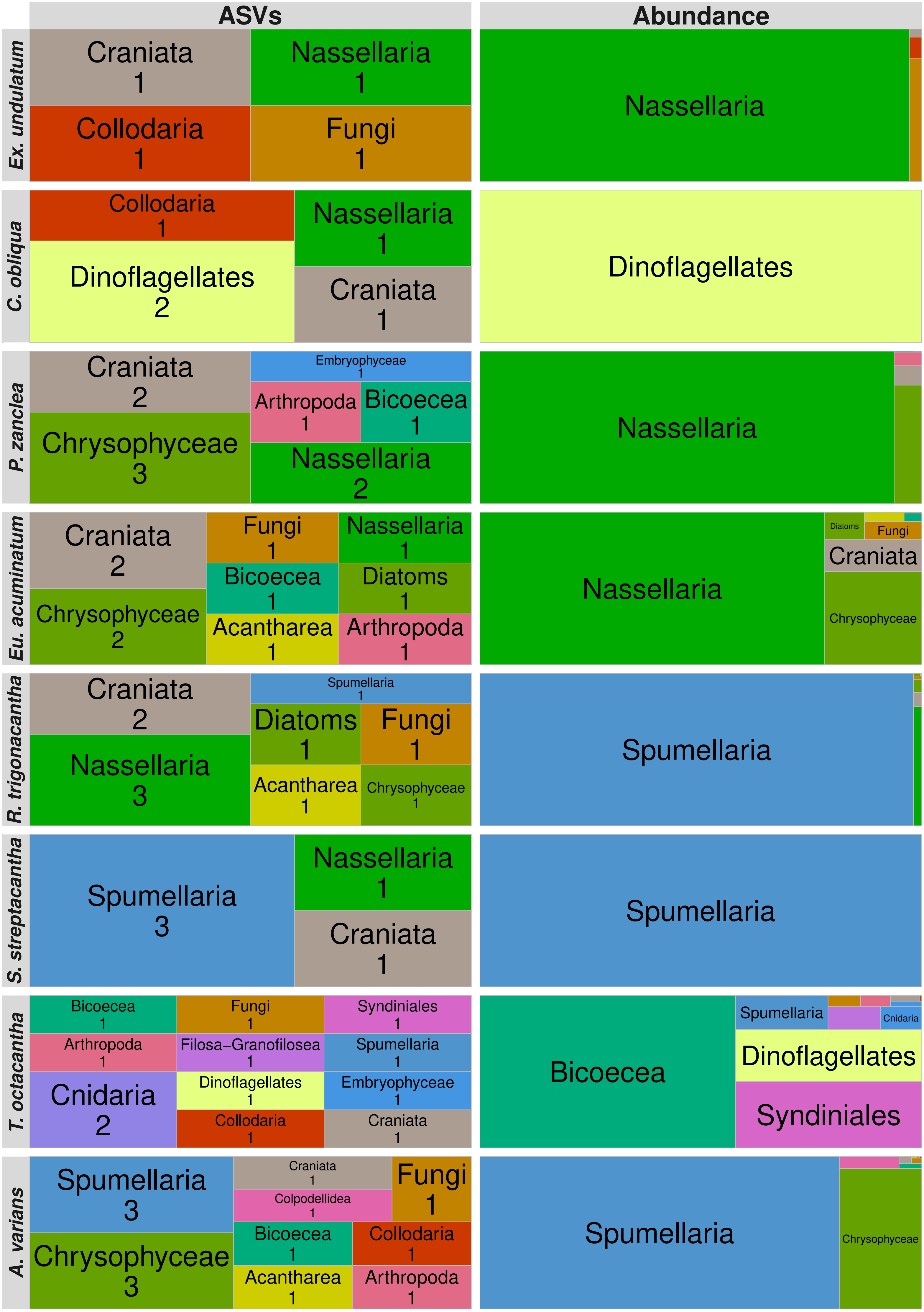

### media-5.pdf

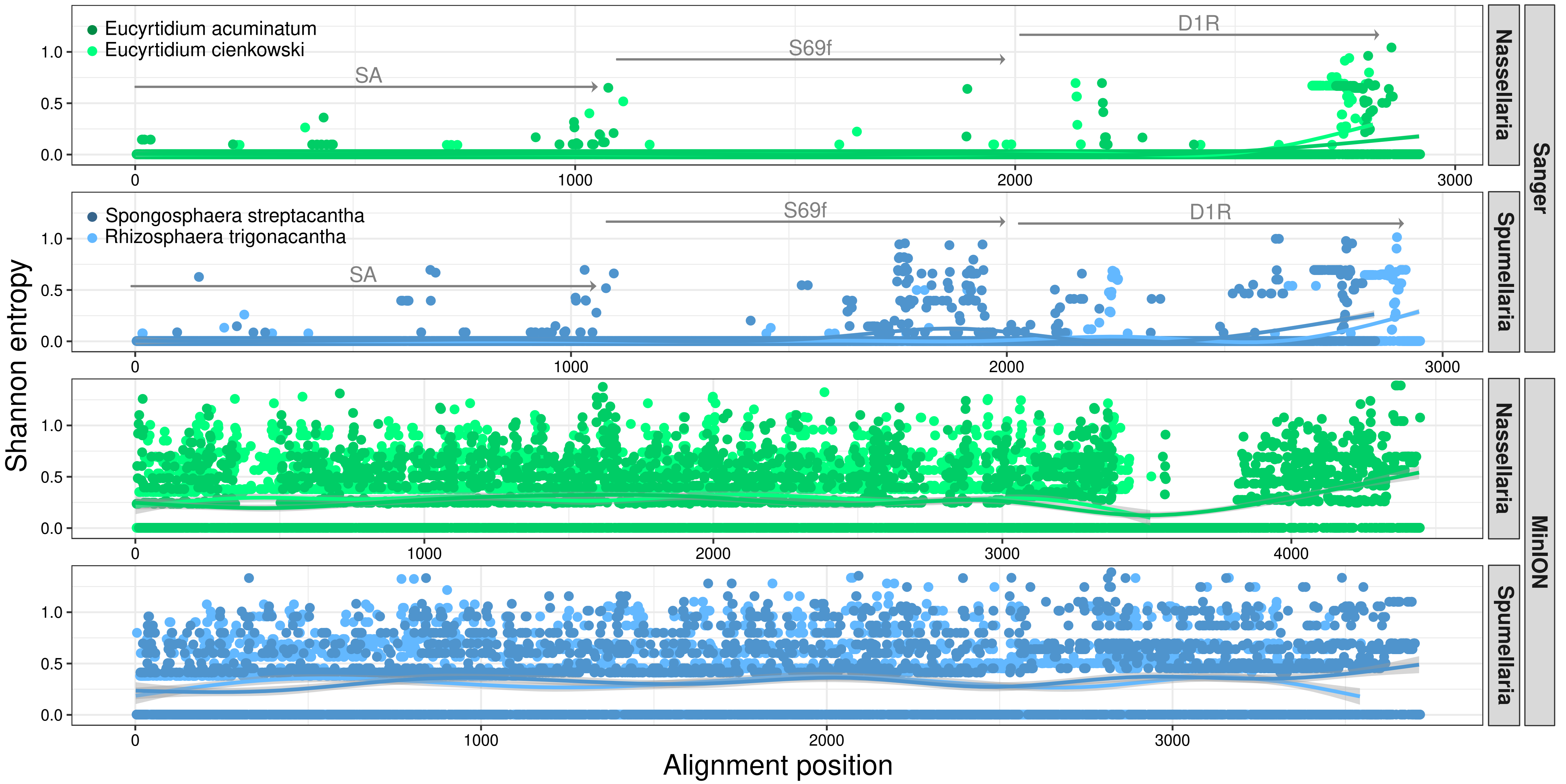
